# A novel and potent MICA/B antibody is therapeutically effective in *KRAS LKB1* mutant lung cancer models

**DOI:** 10.1101/2024.07.30.605880

**Authors:** Ryan R Kowash, Manoj Sabnani, Laura T Gray, Qing Deng, Luc Girard, Yujiro Naito, Kentaro Masuhiro, John D Minna, David E Gerber, Shohei Koyama, Zhiqian Lucy Liu, Hemanta Baruah, Esra A Akbay

## Abstract

Concurrent *KRAS LKB1* (STK11, KL) mutant Non-Small Cell Lung Cancers (NSCLC) is particularly difficult to treat and does not respond well to current immune checkpoint blockade (ICB) therapies. This is due to numerous mechanisms including low antigen presentation limiting T cell mediated killing. To activate anti-tumor immunity, we targeted tumor cell – natural killer (NK) cell interactions. We tested whether a novel antibody based therapeutic strategy that predominantly activates natural killer (NK) cells demonstrates efficacy in pre-clinical mouse models of KL NSCLC. NK cells rely on binding of ligands, such as Major Histocompatibility Complex (MHC) class I-related chain A or B (MICA/B), to the activating receptor NKG2D. Importantly MICA and MICB are widely expressed in elevated levels across NSCLC subtypes including KL lung cancers. Proteases with the tumor microenvironment (TME) can cleave these proteins rendering tumor cells less visible to NK cells. We therefore developed a MICA monoclonal antibody, AHA-1031, which utilizes two NK cell activating receptors. AHA1031 prevents ligand shedding without interfering with binding to NKG2D while targeting cancer cells to antibody mediated cell dependent cytotoxicity (ADCC). Our therapeutic novel antibody has significant monotherapy activity in KL cancer models including xenografts of human cell lines and patient derived xenografts. Activating NK cells through MICA/B stabilization and inducing ADCC offers an alternative and potent therapy option in KL tumors. MICA/B are shed across different tumors making this therapeutic strategy universally applicable.

## Background

Natural killer (NK) cells play a key role in the host innate immune response protecting from pathogens and cancer. NK cells are particularly important for preventing lung cancer as depletion of NK cells in genetically engineered models of lung adenocarcinoma were shown to increase tumor formation (1). Unlike T cells which require MHC for tumor cell recognition and cytotoxic effect, NK cells rely on a complex interplay of both inhibitory and activating ligand-receptor interactions (2). In humans NK cell activating ligands include MICA/B and the UL16 binding proteins (ULBP1-6) (3). Upon their engagement to the NKG2D activating receptor, expressed on both NK cells and T cell subsets, this results in activation of these immune cell populations (2). However, there are numerous mechanisms that tumor cells utilize to avoid immune detection. One of these such mechanisms is the shedding of MICA/B ligands which limits NK and T cell binding to tumor cells leading to decreased immune cell activation (4). Therefore, preventing the shedding of these ligands to target cancer cells to antibody mediated cell dependent cytotoxicity (ADCC) has recently garnered attention as a therapeutic strategy (5).

MICA/B are stress induced ligands and are highly expressed in numerous cancer types including melanoma, glioma, and non-small cell lung cancer (NSCLC) (6-8). However, proteases within the tumor microenvironment can cleave these ligands resulting in reduced expression on tumor cells (9, 10). Soluble MICA/B in the blood has been detected in patients with various cancer types and serum levels are thought to correlate with disease progression and this was demonstrated in NSCLC patients (11)(6, 12). NSCLC is the most common form of lung cancer accounting for around 85% of all lung cancer cases and affecting hundreds of thousands of patients in the United States every year. Immune checkpoint blockade is the current standard of care for NSCLC. However, response rates for some subsets of lung cancer remain low and resistance to current therapies remains a major challenge (13). Therefore, the development of new targeted therapies that overcome current limitations are needed.

The mutational landscape of NSCLC comprises common cancer drivers such as loss of tumor suppressors p53 and LKB1 and activation of oncogenes such as KRAS and EGFR (Epidermal Growth Factor Receptor) (14, 15). LKB1 is encoded by the gene *STK11* and is a key tumor suppressor (16). Concurrent KRAS and LKB1 mutant (KL) NSCLCs are especially difficult to treat and do not respond well to ICB therapy (17). Recent data from a clinical trial testing the efficacy of PD-1 (programmed death 1) (programmed death 1) blockade revealed that K (KRAS) driven tumors had an objective response rate (ORR) of 28.6%, KP (KRAS p53) mutant tumors had an ORR of 35.7%, and KL mutant tumors a dismal ORR of 7.4% (18) underlying the critical need to develop better therapies for this subset of lung cancers.

To develop an alternative immune activating mechanism in ICB resistant tumors, we developed AHA-1031, a novel antibody that targets MICA/B ligands resulting in a potent effect on natural killer cell activation. AHA-1031 targets the α3 domain of MICA/B to prevent their proteolytic cleavage, thereby increasing ligand presence on the cell surface. This facilitates the interaction with the natural killer group 2D (NKG2D) receptor found on NK cells, as well as a subset of T cells including γδ T cells, NKT cells, and CD8+ T cells. We have also incorporated an engineered human IgG1 Fc region to enhance antibody-dependent cell-mediated cytotoxicity (ADCC), which is crucial for NK cell cytotoxicity. Notably, our study is the first to evaluate NK activating therapy agents specifically in KL mutant NSCLC, revealing that effectively engaging multiple potent NK activating receptors enhances NK cell cytotoxicity synergistically. This approach may offer a potential treatment for KL mutant NSCLC without the safety concerns associated with T cell engaging therapies. We focused our study on KL mutant NSCLC tumors due to current treatment ineffectiveness, yet our method is potentially widely applicable to other subtypes of NSCLC or other cancers as MICA/B are widely expressed.

## Materials and Methods

### Expression analysis

NSCLC cell line (19) and patient tumor data (20) were downloaded and analyzed after normalization. Proteomics data was from a previously published study (21). Processed data was downloaded and normalized NKG2DL levels were graphed.

### Flow cytometry

Cells were stained with fixative live dead cell stain (Thermo Fisher Scientific, catalog no. 50-112-1528) for 8 minutes at room temperature. Cells were then stained with fluorophore conjugated antibodies in FACS buffer (2% FBS in PBS) for 20 minutes on ice. Mouse samples were digested for 30 minutes at 37C with 100 units/mL collagenase, Thermo Fisher Scientific, catalogue no. 17104019 10ug/mL DNAase I Sigma catalogue number, 10% heat inactivated FBS in RPMI (Roswell Park Memorial Institute) to disassociate cells. Red blood cells were lysed with ACK lysis buffer (Thermo Fisher Scientific, catalogue no A1049201). Tissues were passed through 70um cell strainer to make single cell suspension. Cells were stained with fixative live dead cell stain followed by CD16/32 antibody (BioLegend, catalog no. 101320) for 20 minutes on ice. Next, they were incubated with fluorophore-conjugated antibodies. If applicable, intracellular staining was performed by eBioscience Foxp3/Transcription Factor Staining Buffer Set (Thermo Fisher Scientific, catalog no. 00-5523-00) according to the manufacturer’s instructions. To determine T-cell and NK cell activation, lymphocytes were enriched using Ficoll-Paque (GE Healthcare) following the protocol. Enriched samples were incubated with PMA/ionomycin/Golgi plug for ex vivo stimulation for 6 hours. The same staining protocol as detailed above was utilized. Samples were run on BDFACS Canto, and flow data were analyzed using FlowJo. Source and catalog numbers of all antibodies used for flow are outlined in Supplementary Table 1.

### Antibody production

Production of recombinant antibodies was conducted either in EXPI293 cells or outsourced to Wuxi Biologics, where it was produced in CHO-K cells. The process involved utilizing two plasmids, one encoding the light chain (pcDNA 3.1 LC) and the other encoding the heavy chain (pcDNA 3.1 HC). For EXPI293 production, these plasmids were co-transfected into Expi293F suspension cells using lipid based ExpiFectamine 293 transfection kit (catalog no. A14525, Thermo Fisher Scientific). All procedures followed the guidelines outlined in the Life Technologies ExpiFectamine 293 transfection kit. Transfected cells were cultured at 37 °C with 8% CO2 and 120 rpm. After 5-6 days post-transfection, supernatants underwent protein affinity purification using a high-performance 5 mL HiTrap MabSelect SuRe column (Cytiva, catalog no. 11003494) from GE Healthcare Life Sciences for large-scale production. Production in CHO-K cells was outsourced to Wuxi Biologics in China.

### Shedding inhibition of MICA on cancer cell surface by alpha3 domain specific MICA antibodies (ELISA and Flow based Assays)

Human cell lines A375 (CRL-1619), A549 (CCL-185), H2030 (CRL-5914), NCI-H226 (CRL-5826), Calu3 (HTB-55), PANC1 (CRL-169), Capan2 (HTB-80), OVACAR3 (HTB-161), HCT116, 22Rv1 (CRL-2505), PC3 (CRL-1435), and LNCaP (CRL-1740) were obtained from the American Type Culture Collection (ATCC, USA). All cell lines were cultured in their recommended media with 1% antibiotic mixture (Penicillin/Streptomycin – Pen/Strep, 10,000 IU/mL; Sigma-Aldrich) and 10% Fetal Bovine Serum (FBS; Gibco, Thermo Fisher Scientific). The cells were maintained at 37°C in a humidified atmosphere containing 5% carbon dioxide (CO2).

Tumor cells were cultured with our α3 domain specific anti-MICA antibodies and control benchmark antibodies (including Hu Isotype IgG1 and cell only group) for 24 to 72 hours in cell-culture treated 96-well plates. The antibodies were titrated from 200 nM down to sub-picomolar concentrations. Supernatants were harvested and MICA levels were quantified via sandwich ELISA (R&D Systems, DUO Set Human MICA DY 1300). Cells were detached using Versene (Gibco, catalog no. 15040-066) to quantify cell surface MICA stabilization then spun down and washed with 1X PBS and stained with fixative live dead cell stain (Invitrogen eBioscience, 65-0865-14). Cells were washed and stained with a PE-conjugated anti-MICA antibody (Clone 6D4, Biolegend, 320902) in FACS buffer (2% FBS in PBS) for 20 minutes on ice. Samples were analyzed using flow cytometer Attune™NxT (Thermo Scientific) and data was analyzed with FlowJo software.

### Isolation of NK cells from donor PBMCs (Peripheral blood mononuclear cells)

Peripheral blood mononuclear cells (PBMCs) were isolated via Ficoll-Paque gradient centrifugation, employing products from Amersham Pharmacia Biotech (Uppsala, Sweden), using buffy coat obtained from Carter BloodCare (Bedford, TX). Human primary NK cells were isolated from the cryopreserved PBMCs utilizing the NK Cell Isolation Kit (Miltenyi Biotec, catalog no. 130-092-657) according to the manufacturer’s instructions.

### Antibody dependent cytotoxicity assays

The antibody-dependent cellular cytotoxicity (ADCC) activity of human primary NK cells was evaluated using a two-color flow cytometric assay (22). Target tumor cells (A375/A549; 1 × 10^6^) were incubated with 1 μl of 0.5 mM carboxyfluorescein diacetate succinimidyl ester (CFSE; Invitrogen, Eugene, OR, USA) in phosphate-buffered saline (PBS) for 10 min at 37°C. The cells were washed thrice with ice-cold RPMI 1640 medium. Effector NK cells were mixed with the target cells at an effector-to-target (E/T) ratio of 10:1 (5.0 × 10^6^/ml effector cells, 5 × 10^5^/ml target cells; final volume: 200 μl) and incubated with or without antibodies (AHA-1032, AHA-1031, Human Isotype control) at a concentration of 20 nM, titrated down to sub-picomolar levels overnight. Cells were washed twice with 1X PBS and stained with 7-aminoactinomycin D (7-AAD; eBioscience, San Diego, CA, USA) for subsequent analysis via two-color flow cytometry, conducted on an Attune™ device (Thermo Scientific). CFSE and 7AAD double-positive cells were indicative of tumor target cells lysed via ADCC. The EC50 was calculated using nonlinear regression analysis performed with GraphPad Prism.

### MICA ELISA of patient blood and cell lines

A549, H2030, and H69 cells were plated in 6-well plate at density of 0.5 million/well. Cells were cultured in complete growth medium and cell culture media was collected 48 hours later. Culture media was centrifuged at 1500 rpm for 10 min at 4°C to collect the supernatant. The analysis of MICA concentration in cell culture supernatant was conducted according to the instructions of Human MICA ELISA kit (R&D systems, catalog no. DY1300).

Tumor volumes were calculated based on the sum of the diameters of target lesions, including primary tumors and metastatic tumors, according to the RECIST criteria (version 1.1).

The objective response was evaluated at 6 weeks after treatment and was repeated every 6 weeks thereafter. Responders (= sensitive) were defined as PR (partial response) or SD (stable disease), and non-responders (= resistance) as PD (progressive disease) at 3 months (6 cycles x 2 weeks) after initiation of Nivolumab treatment. All patient samples were obtained from subjects providing written informed consent for blood in accordance with the Declaration of Helsinki and studies were approved by an Institutional review board at Osaka University (Suita, Japan; no.11122, 16450). MICA levels were quantified as above.

### Mouse xenograft and PDX models

All mice experiments were performed following Institutional Animal Care and Use Committee (IACUC) and ARRIVE guidelines. Human cell lines were implanted into athymic nude mice (The Jackson Laboratory, catalog no. 002019) of 4–8 weeks of age. Mice were injected subcutaneously with 5 million A549 or H2030 human cells with 50% total volume matrigel (Corning, catalog no. 354230). A KRAS-LKB1 (KL) PDX NSCLC model was obtained from the Hamon Cancer Center at UTSW (UT Southwestern), and tumor pieces were implanted into the flanks of nude mice. Vehicle (PBS) or 10mg/kg AHA-1031 were injected by IP. Volume (mm^3^) = width (mm) × width (mm) × length (mm)/2. Male and female mice were both used, evenly distributed, and randomized for treatment and started when tumors reached 75-100mm^3^.

### Cell culture

Human cell Lines A549, H69, and H2030 were grown in RPMI1640 media supplemented with 10% FBS and 1% antibiotic/antimytotic. All cell lines were tested for mycoplasma and confirmed negative.

### Statistical analysis

All statistical analysis was completed with Graphpad prism software. *P* values were determined using single or multiple unpaired T tests for analysis. Significant *P* values are shown as: *: p < 0.05, ^∗∗^: p < 0.01, ^∗∗∗^: p < 0.001, ^∗∗∗∗^: p < 0.0001.

### Data availability statement

Publicly available datasets were used for the analysis of gene expression and proteomics.

Data from antibody screen is available from the corresponding authors upon reasonable request.

## Results

### NSCLCs express and shed MICA and MICB proteins

We analyzed expression of all NKG2DLs MICA, MICB, and ULBP1-6 in lung adenocarcinoma cell lines and primary patient tissue from the lung adenocarcinoma samples from the lung cancer genome atlas (TCGA) data set (19, 20). MICA/B and ULBP1-3 are highly expressed while ULBP4-6 are expressed at lower levels. MICA is the highest expressed NKG2D ligand in lung adenocarcinomas both for cell lines and tumor tissue at the RNA level warranting further investigation (Figure 1A and 1B). Importantly, analysis of NKG2DL expression specifically in KL mutant tumors showed that MICA is expressed at the highest level (Supplemental Figure 1A). Previous reports showed that KL mutant tumors have decreased gene expression of key immune related genes including HLA (Human Leukocyte Antigen) related genes and TAP1 (23). We had previously shown that lung adenocarcinomas also shed these ligands (7). We validated this in a large cohort of patient samples composed of 1400 blood samples from patients with common cancer types including lung cancer, breast cancer, colorectal cancer, ovarian cancer, endometrial cancer, prostate cancer, glioma, and hematological cancers-myeloma, acute myeloid leukemia (AML), chronic lymphocytic leukemia (CLL), and lymphoma. (Figure 1C). This dataset included blood samples from 268 lung cancer patients. In this proteomics-based assay MICA/B was elevated in the blood of majority of patients from all cancers in this dataset (21).

**Figure 1:**
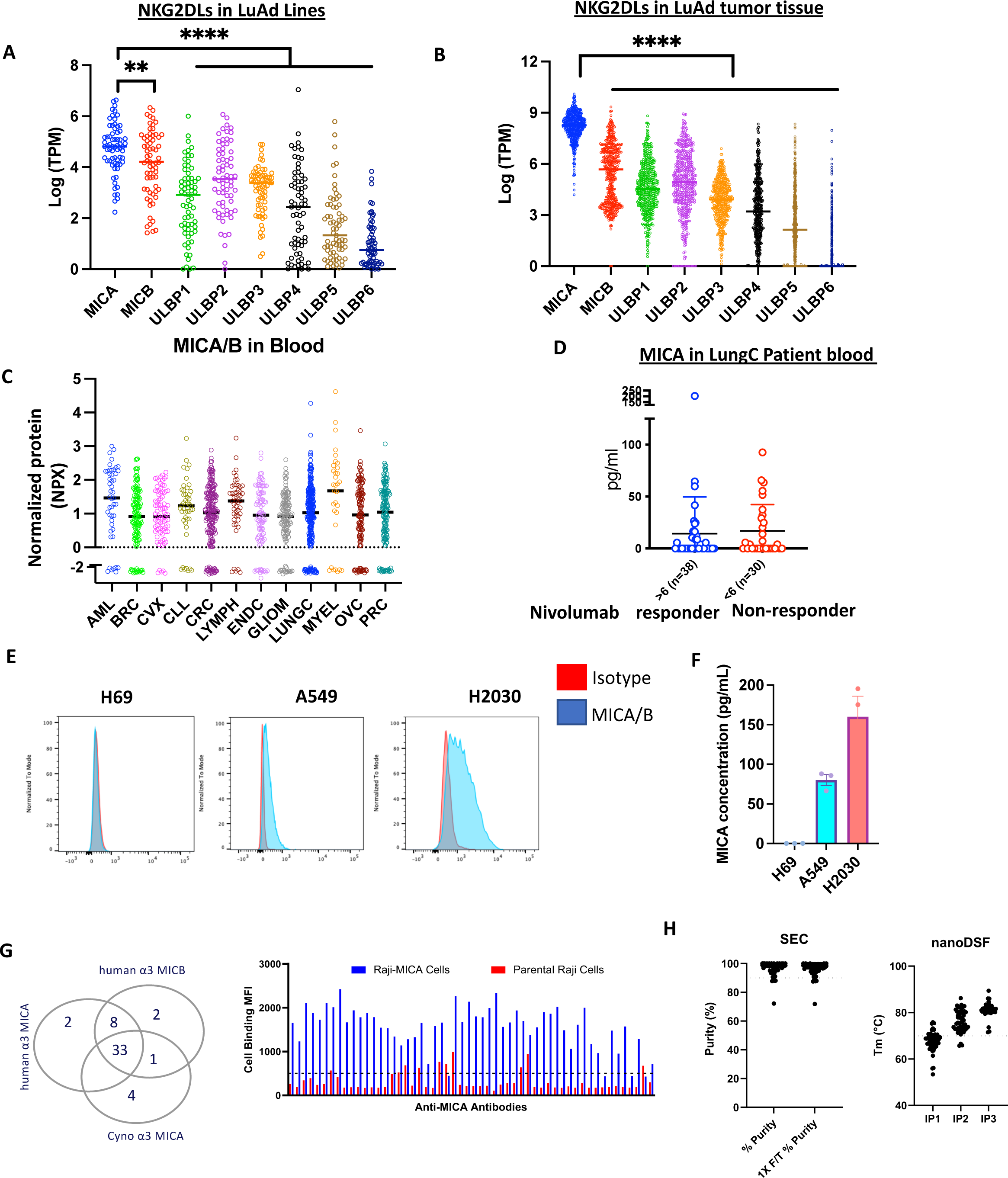
MICA/B expression and shedding from cancers is a targetable feature. (A) mRNA expression displayed as normalized Log (TPM) of NK cell ligands MICA, MICB, ULBP1, ULBP2, and ULBP3 in lung adenocarcinoma cell lines (n=67) (B) mRNA expression of NK cell ligands MICA, MICB, ULBP1, ULBP2, and ULBP3 in lung adenocarcinoma patient tumors (n=490) (C) Normalized MICA/B levels in the blood of cancer patients determined by proximity ligation assay (D) ELISA analysis of secreted MICA in the blood of NSCLC patients that were sensitive (n=38) or resistant (n=30) to Nivolumab treatment (E) Cell surface expression of MICA/B in human SCLC cell line H69 and human NSCLC cell lines A549 and H2030 as determined by flow cytometry (F) ELISA analysis of MICA secretion (pg/mL) in SCLC cell line H69 as well as KL mutant NSCLC cell lines A549 and H2030 (G) Screening of alpha3 specific antibodies cross reactivities against human MICA/MICB and cynomolgus MICA. Binding of alpha3 MICA unique clones against RAJI parental and RAJI MICA transduced cell line (H) Physicochemical properties for AHA-1031. a-% of main peak by SEC after protein A purification and % of main peak by SEC after1freeze/thaw cycle. f-nano-Differential scanning fluorimetry, or nanoDSF, is a biophysical characterization technique used for assessing the conformational stability of a biological sample. **, P<0.01, ****, P<0.001.

We next validated this in an independent cohort of patient samples where we had information about patients’ response to ICB antibody-Nivolumab. Consistent with the large dataset, NSCLC patients had MICA in their blood as determined by ELISA (Figure 1D). Response to Nivolumab was determined based on RECIST criteria for primary tumors and metastatic tumors. The objective response was evaluated at 6 weeks after treatment and was repeated every 6 weeks thereafter. Responders (= sensitive) were defined as Partial response or Stable disease, and non-responders (= resistance) as progressive disease at 3 months (6 cycles every 2 weeks) after initiation of Nivolumab treatment. There were no significant differences between patients whose tumors responded and patients whose tumors did not respond to Nivolumab indicating that MICA targeting can be considered for all these patients (Supplemental Figure 1B).

To determine expression levels of MICA/B specifically in the models used in this study, we first confirmed tumor cell surface expression KL mutant NSCLC A549 and H2030 by flow cytometry. Consistent with RNA data, these cells expressed elevated levels of MICA/B. A small-cell lung cancer (SCLC) cell line H69 did not express surface MICA/B as expected (Figure 1E). These data align with previous reports that NSCLCs can be targets for NK cells (7, 24). We next wanted to determine whether these cell lines shed MICA in the media in cell culture by ELISA. We observed that levels of soluble MICA in the media correlated with their cell surface expression and that KL lines shed MICA/B into the media whereas as SCLC cell line H69 with no surface MICA/B expression did not (Figure 1F).

After establishing that MICA is a putative cancer target, we next identified high affinity and cross-reactive antibodies to cynomolgus (cyno) MICA. Dissociation constant (KD) measurements were performed for all the 54 purified antibodies by surface plasmon resonance (SPR) against human full-length MICA, MICB, the alpha 3 domain of MICA and full-length MICA. Our three-prong discovery effort was productive. Briefly, all 54 clones bind to full length human MICA, 41 clones bind to the desired alpha 3 domain of both human MICA and MICB, 33 of those 41 mAbs cross react with the alpha 3 domain of cynomolgus MICA (Figure 1G and Supplementary Figure 2). We next performed cell binding of the entire panel of 54 antibodies to confirm that the antibodies that bind to recombinant antigens are also able to bind MICA expressed on the cell surface. More than 80% of the antibodies bind specifically to MICA-overexpressing Raji cells compared to parental Raji cells (Figure 1G). We also did extensive binning analysis by SPR and identified as many as 6 different binning epitopes. This diversity in binding epitope allowed us to identify multiple mAbs with the right biology and developability profiles. SEC profiles for all 54 purified antibodies were acquired before and after freeze thaw. Greater than 90% of the antibodies showed a >90% monomeric peak, suggesting robust homogeneity and stability (data not shown). The thermal stability of the antibodies as assessed via nano differential scanning fluorimetry (nanoDSF) were in a similar range to that of clinical antibodies (Figure 1H).

### Development of a shedding preventing MICA/B specific antibody

Our first-generation lead candidate identified with robust cyno cross reactivity and clean developability profile is named AHA-1032. All 4 top candidates have the desired properties to be the lead candidates, such as high affinity binding to human MICA or MICB, and binding to the desired alpha 3 domain of MICA, and more importantly, can kill cancer cells with high potency. AHA-1032 has an additional advantage in that it binds robustly to cyno MICA with similar monovalent affinity as human MICA or MICB (Figure 2A). This excellent cross reactivity allows for accurate translation of pharmacokinetics (PK)/pharmacodynamics (PD) data from IND (Investigational New Drug) enabling cyno toxicity studies into clinical trials. Moreover, when we compare the developability profiles of the four antibodies, AHA-1032 has the most optimal profile with no weaknesses in any of the assays we performed (Supplementary figure 2). The EC_50_ binding value of AHA-1032 (MICA α3 specific) to MICA overexpressing Raji cells was determined and found to correlate well with recombinant MICA binding (Figure 2B).

**Figure 2:**
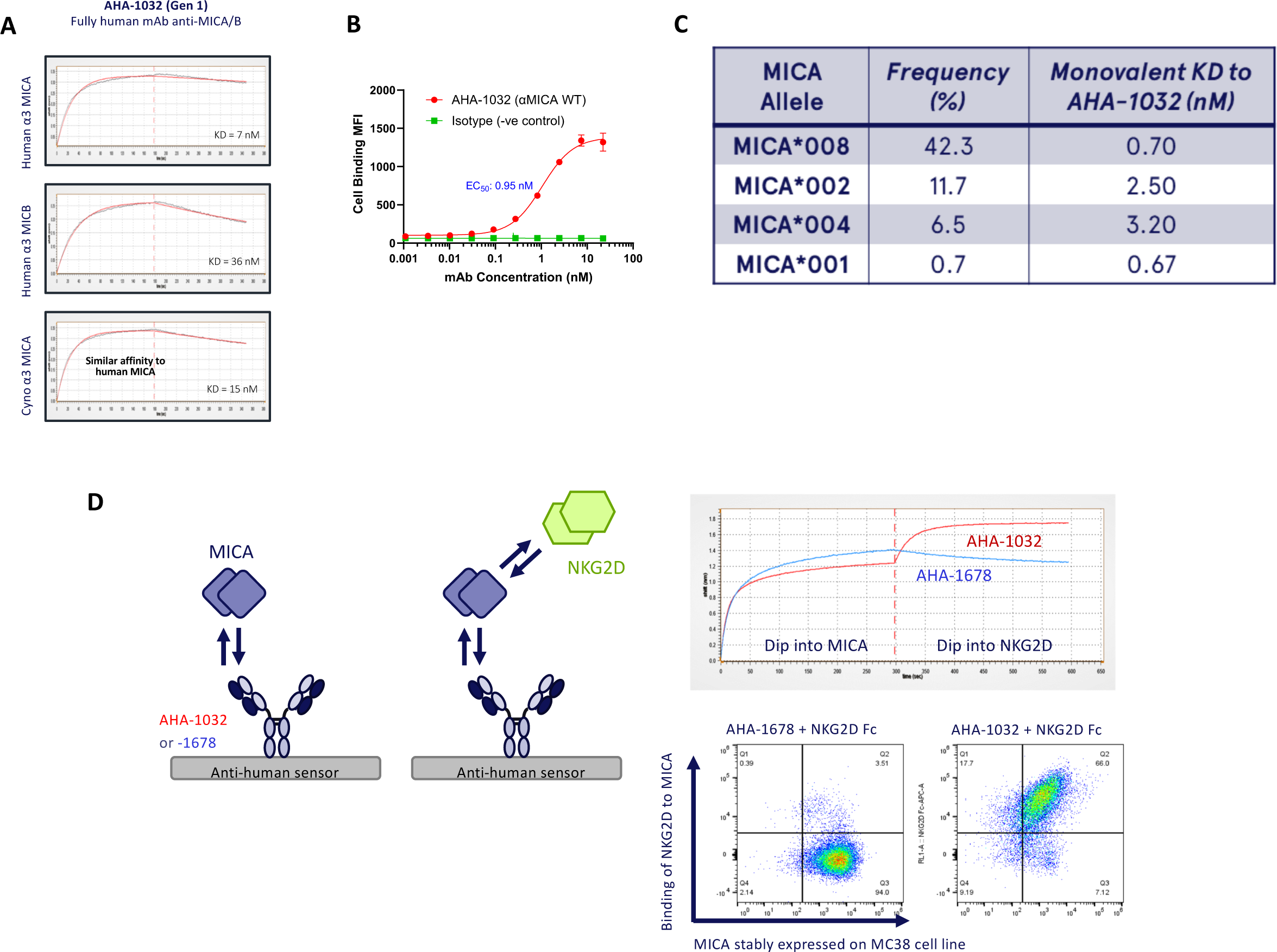
Binding characterization and MICA shedding inhibition of AHA-1031 and 1032. A) Binding of AHA-1032 with α3 specific Human MICA/MICB and Cynomolgus MICA B) α3 specific anti MICA/MICB binding on Bio layer Interferometry (BLI system Gator Bio) and binding with Parental RAJI cells and MICA transduced cells. C) MICA isoform frequency and KD of AHA-1032 D) Binding of AHA-1032 and NKG2D receptor, Binding of 1032 and 1031 (MICA α3 specific) is independent of NKG2D binding.

One other complexity about MICA is the polymorphic nature of the protein. There are more than 100 identified alleles and developing an antibody that binds to most of the alleles is an important criterion to maximize patient coverage (25). Notably, there are a few variants such MICA*008 (42.3%) that are more frequently observed in cancer patient populations. We therefore cloned four such MICA variants and confirmed binding to AHA-1032. This gives a total coverage of as much as 61.2% of all the variants. Next, we compared the sequence homology of the alpha 3 domain of these four variants to the other remaining variants and found that they are greater than 95% similar, indicating that AHA-1032 would likely bind to most of the variants (Figure 2C).

Additionally, we performed multiple developability assays including binding to baculovirus particles (BVP), soluble membrane preparations (SMP), single-stranded DNA, insulin, and affinity-capture self-interaction nanoparticle spectroscopy (AC-SINS) (Supplementary Figure 3A). Based on initial data, we selected 26 antibodies for further evaluation of MICA shedding inhibition properties. From these, we identified 4 top candidates and assessed their ability to induce NK cell-mediated killing. Moreover, we conducted an early assessment of the clinical utility potential of the germlines of our antibodies and immunogenicity profiling using two methods: Humanness and T20 scores, comparing them to Nivolumab, discovered using a similar transgenic mouse platform (Ultimab, Medarex). Our lead candidates exhibit features like FDA-approved antibodies, giving confidence for further development (Supplementary figure 3B and 3C). Based on our overall findings, we selected AHA-1032 as our first-generation lead candidate.

Next, we confirmed by biolayer interferometry (BLI) that the binding of AHA-1032 to the membrane proximal α3 MICA domain does not interfere with NKG2D binding to MICA. On the contrary, an antibody which binds to the α1 and α2 MICA domains, such as control AHA-1678, blocks the binding of NKG2D to MICA. We were also able to translate our BLI findings to cellular context by demonstrating simultaneous binding of MICA expressed on MC-38 cells to both AHA-1032 and NKG2D-Fc. The alpha3 specific anti MICA/MICB AHA-1032 binding does not interfere with MICA/MICB – NKG2D binding. The MC38 cell line expressing human MICA was used to prove this interaction. MC38 expressing MICA cells were stained with either alpha 3 specific AHA-1032 or Alpha1/Alpha2 specific mAb AHA-1678, Cells were stained with NKG2D Fc to test monoclonal antibody binding interfering with NKG2D binding (Figure 2D).

Prevention of shedding was confirmed in lung cancer lines H2030, A549, H226, Calu-3. Beyond lung cancer models, we also observe that AHA-1031 stabilizes MICA/B expression in pancreatic (PANC1, Capan-2), colon (HCT116), ovarian (OVCAR3), and prostate cancer cell lines (22Rv1, PC3, LNCaP). This data therefore demonstrates broad activity of AHA1032 across cancer types (Figure 3A and 3B). Given its significantly enhanced cell killing properties, we selected AHA-1031 as our lead candidate. Additionally, when comparing the ability of AHA-1031 and AHA-1032 for specific cancer cell killing mediated by primary NK cells, AHA-1031 exhibited far superior performance (Figure 3C).

**Figure 3:**
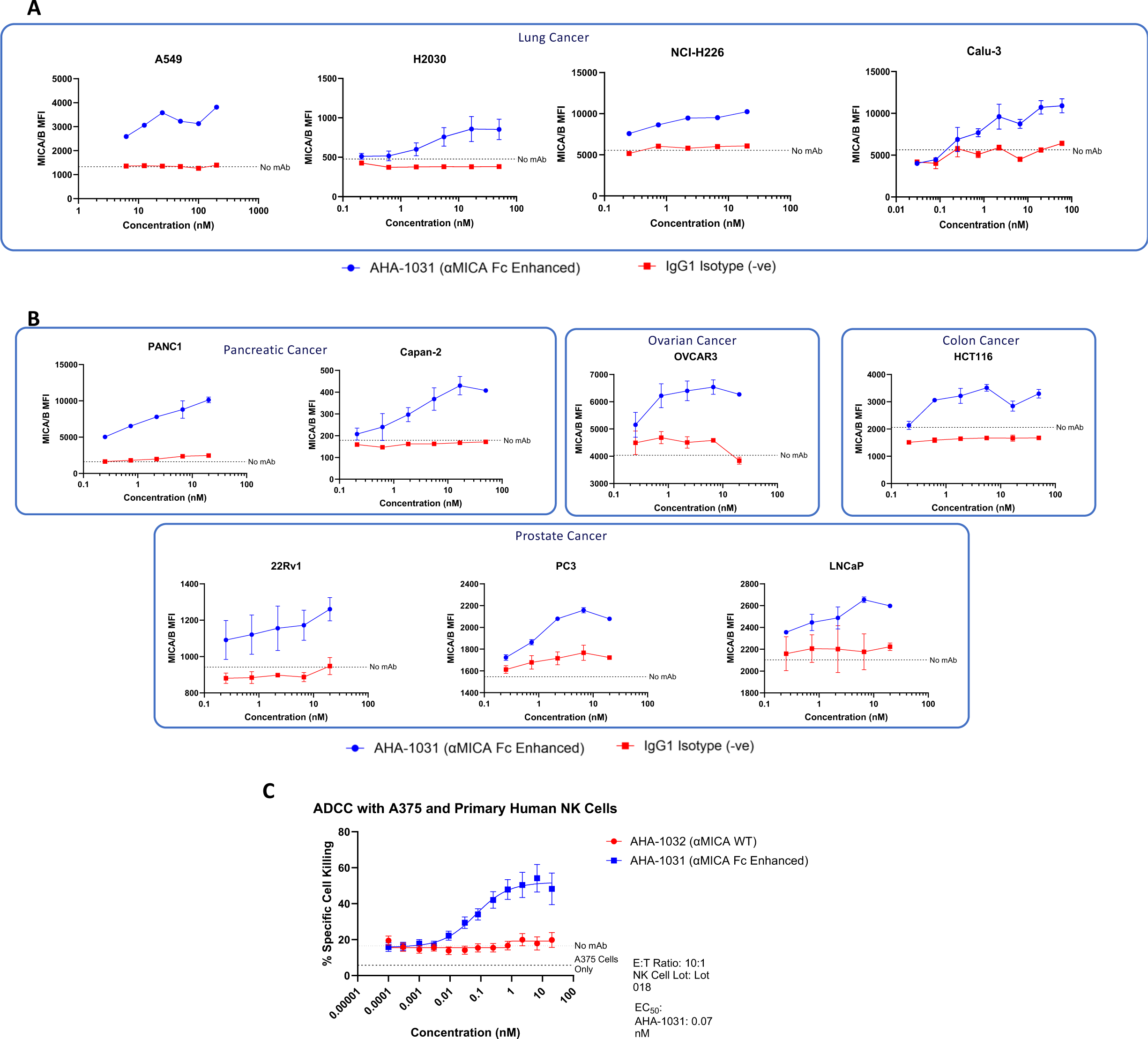
MICA shedding inhibition in cancer cell lines. (A) Stabilization of MICA on the cell surface as shown through flow cytometry in lung cancer cell lines A549, H2030, NCI-H226, and Calu-3 incubated with increasing doses of isotype control or AHA-1031 (B) Stabilization of MICA on the cell surface as shown through flow cytometry in pancreatic, ovarian, colon, and prostate cancer cell lines incubated with increasing doses of isotype control or AHA-1031 (C) ADCC as evaluated by two color assay in A375 cells treated with isolated NK cells from PBMCs and AHA-1031 or AHA-1032.

### Characterization of novel MICA Antibodies AHA-1031 and AHA-1032; engineered IgG1-Fc (AHA-1031) improved cell killing significantly

With production of AHA-1032 we demonstrated specificity and that increasing concentrations of antibody stabilized MICA shedding (Figure 4A). Concurrently, through flow cytometry analysis AHA-1032 stabilized MICA ligand expression on the cell surface (Figure 4B). We next aimed to enhance the efficacy of our first-generation drug candidate in killing cancer cells by boosting ADCC activity. Efficient activation of NK cells relies on the engagement of multiple NK cell receptors, including MICA/NKG2D and CD16a/high-affinity engineered IgG1-Fc. To achieve this, we investigated a modified construct by substituting the wild type IgG1-Fc of AHA-1032 with an amino acid-engineered IgG1-Fc (AHA-1031). In a head-to-head comparison using a CD16+ Jurkat reporter cell assay with NFAT-mediated luciferase activation, AHA-1031, the second-generation construct with engineered IgG1-Fc, demonstrated a 7-fold increase in CD16-mediated NFAT activation compared to AHA-1032 (Figure 4C).

**Figure 4:**
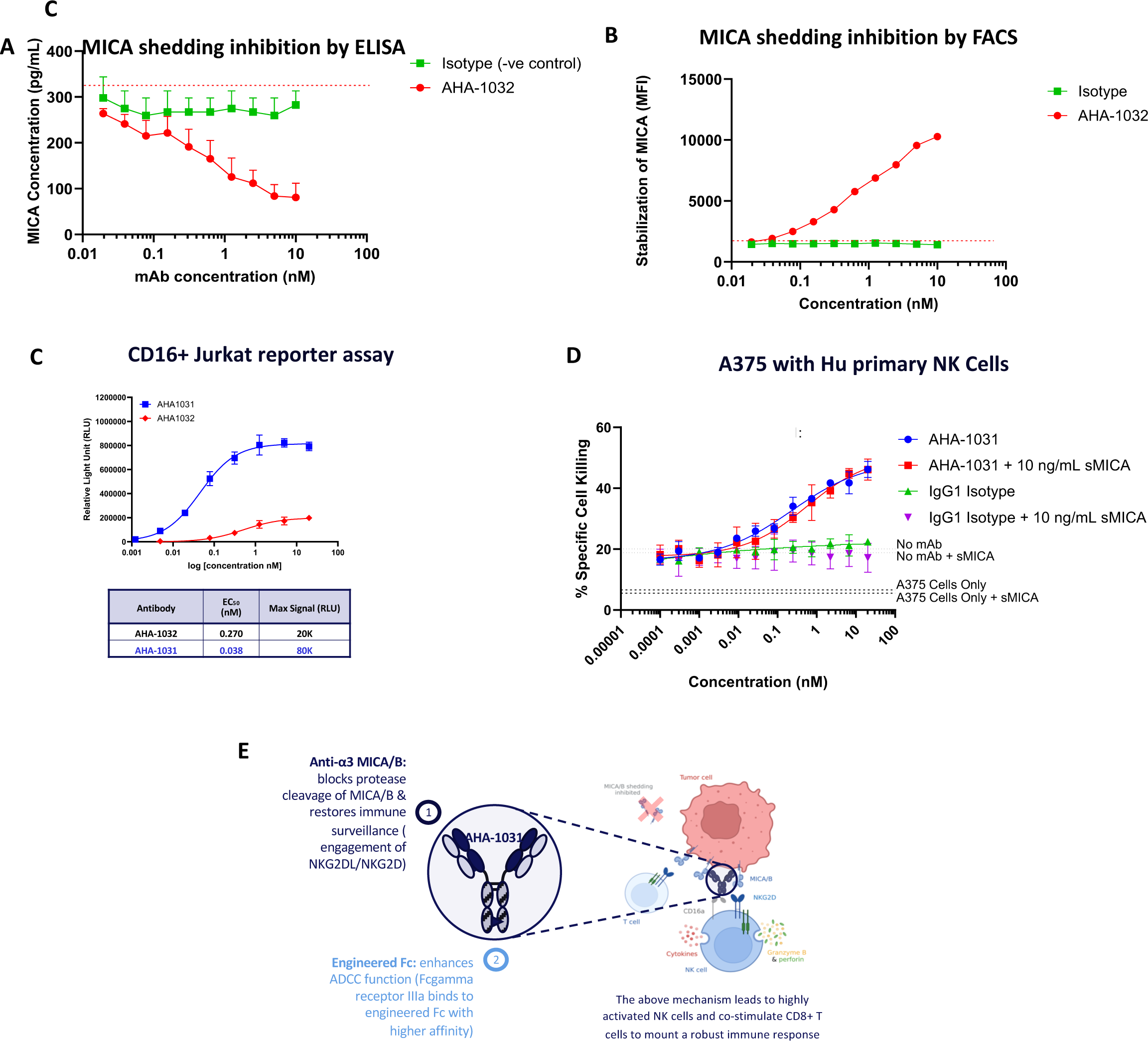
AHA1031 inhibits MICA shedding from cancer cells. (A) MICA shedding inhibition through ELISA analysis of A375 cells incubated with isotype versus AHA-1032 at increasing doses of antibody (B) Flow cytometry analysis of staining of A375 melanoma cells pre-incubated with AHA-1032 using 6D4-PE and stained with an anti-MICA antibody that binds to the α1 and α2 domains. MICA on cell surface with antibody concentration was graphed (C) CD16+ Jurkat reporter cell assay with NFAT-mediated luciferase activation in AHA-1031 vs AHA-1032 displaying EC50 (nM) and max signal (RLU) quantification (D) ADCC activity of A375 cells incubated with primary NK cells in the presence of or absence of shed MICA (E) Aakha’s best-in-class anti-MICA/B antibodies leads to highly activated NK cells and potentially co-stimulate CD8+ T cells to mount a robust immune response via two main antibody features: (1) anti-α3 MICA/B arm: blocks protease cleavage of MICA/B and restores immune surveillance (engagement of NKG2DL/NKG2D), and (2) engineered Fc: enhances ADCC function (Fc gamma receptor IIIa binds to engineered Fc with higher affinity). Illustration made with Biorender

We further established that there is no discernible difference in ADCC activity in the presence of soluble MICA up to a concentration of 10 ng/ml (Figure 4D). To our knowledge, this concentration represents the highest level of shed MICA reported in blood serum for metastatic castrate-resistant prostate cancer (26). In contrast, other shedding antigens such as MUC16, MUC-1, and CEACAM5 are typically shed at significantly higher levels (ug/ml) than MICA. This variance creates a “sink effect,” posing a considerable challenge for antibodies targeting those tumors associated antigen (TAA) to effectively reach the intended tumor site.

Based on this data we propose the following mechanism of action for AHA-1031: It activates NK cells and potentially co-stimulate CD8+ T cells to mount a robust immune response via two main antibody features: (1) anti-α3 MICA/B arm: blocks protease cleavage of MICA/B and restores immune surveillance (engagement of NKG2DL/NKG2D), and (2) engineered Fc: enhances ADCC function (Fc gamma receptor IIIa binds to engineered Fc with higher affinity). The Anti-alpha3 MICA/B arm blocks protease cleavage of MICA/B, thereby restoring immune surveillance by facilitating the engagement of NKG2DL/NKG2D receptors (Figure 4E). Additionally, the engineered Fc region enhances ADCC function by promoting stronger binding to Fc gamma receptor IIIa, thereby contribute to the potent immune response elicited by AHA-1031, making it a promising candidate for cancer immunotherapy.

### Therapeutic efficacy and immune mediated effects of AHA 1031 in *KRAS LKB1* mutant xenograft models

With our in vitro results demonstrating anti-tumor activity and prevention of MICA shedding, we next evaluated the efficacy of our antibody in vivo. Given that the mouse NKG2D and FcεR receptors can bind functionally to human NKG2DLs and IgG1 respectively, we evaluated in vivo efficacy of AHA-1031 in nude mice with MICA expressing human lung cancer cells. In H2030 xenografts we observed significantly decreased tumor growth with AHA-1032 which has a WT engineered Fc region (Figure 5A). We observed further improved efficacy with AHA-1031 with our ADDC enhanced therapeutic. Importantly, in our AHA-1032 treated animals we observed no complete tumor response whereas with AHA-1031 2 out of 6 animals were completely cleared of tumor (Figure 5B). Importantly we detected no observable toxicity of our therapeutic through measurement of mouse bodyweight during and after treatment (Supplemental Figure 4A and 4B). Next, we evaluated the levels of the liver enzymes alanine transaminase (ALT) and aspartate transferase (AST), as well as levels of the cytokines IFNγ, TNFα, and IL-2, in the serum of B16F10-MICA tumor-bearing mice after repeated dosing of AHA-1031. We did not observe an increase in any of these toxicity markers over levels observed in PBS-treated mice (Supplementary Figure 4C). Our results demonstrate superior efficacy in this experiment of our AHA-1031 therapeutic likely due to increased ADCC activity. Importantly a recent study demonstrated that the toxicity of NK cells is enhanced in PD-L1 (programmed death ligand 1) positive tumor models with the addition of ADCC activity (27). This therefore confirms an important aspect of our design of AHA-1031 with an engineered ADCC Fc region.

**Figure 5:**
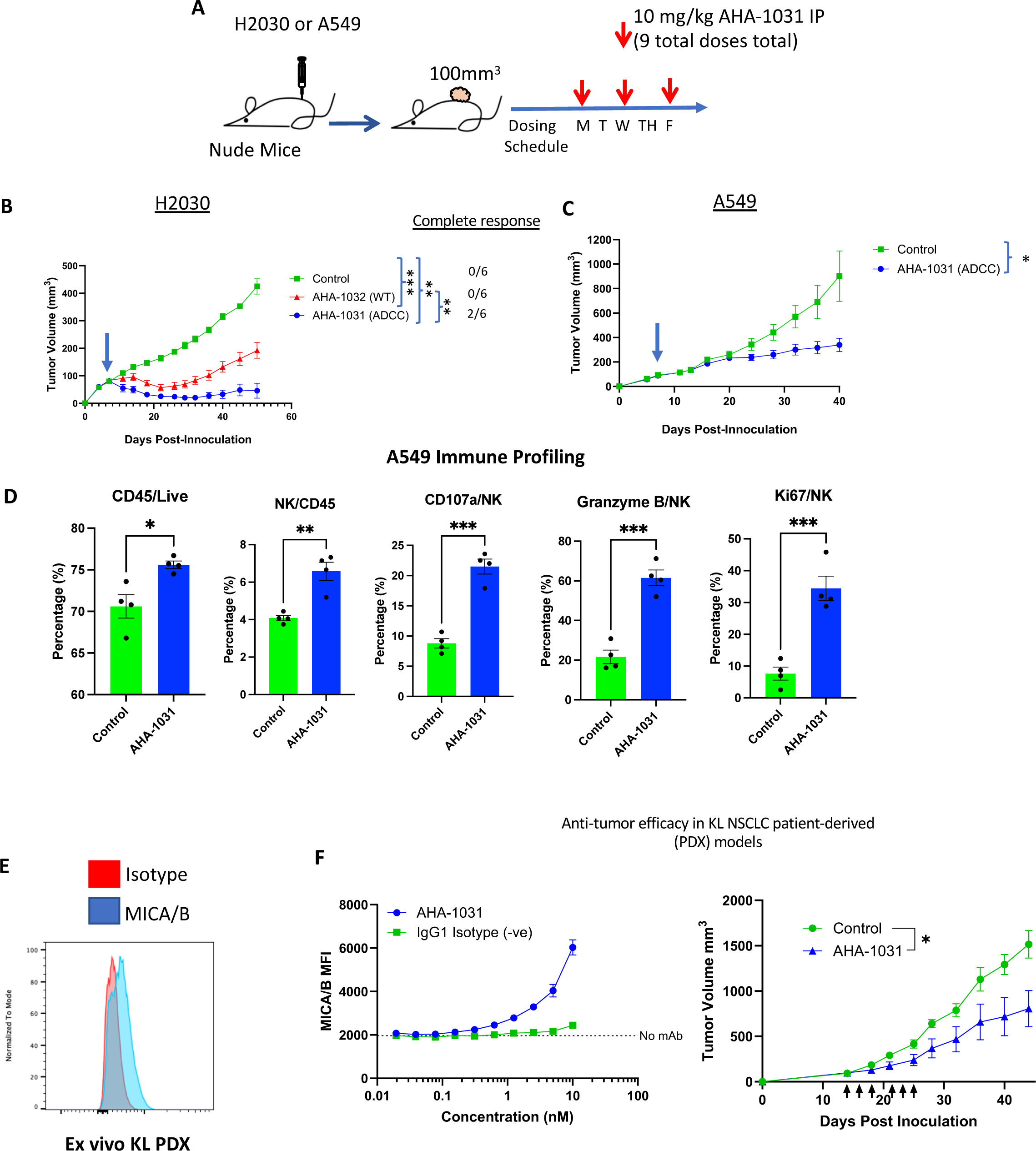
Efficacy of AHA in vitro and in vivo. (A) Schema displaying injection of 5 million A549 or H2030 subcutaneously into nude outbred mice. Tumors were monitored and when tumor volume reached 100mm^3^ mice were started on 10mg/kg AHA-1031 IP MWF for a total of 9 doses (B) Subcutaneous tumor growth curve of H2030 cells injected into nude mice treated with control (n=6), AHA-1032 WT (n=6), or AHA-1031 ADCC (n=6). The number of complete responders is also shown for each group (C) Subcutaneous tumor growth curve of A549 cells injected into nude mice treated with vehicle (n=6) or AHA-1031 (n=6) (D) Flow cytometry analysis of subcutaneous A549 tumor tissue from control (n=4) or AHA-1031 (n=4) treated mice. Immune cell markers for CD45/live, NK/CD45, CD107a/NK, Granzyme B/NK, and Ki67/NK were measured n=4 for control and AHA-1031 (E) Flow cytometry staining of MICA/B in KL PDX ex vivo sample (F) Ex vivo AHA-1031 stabilization of MICA on KL PDX tumor cells determined by flow cytometry (left). Tumor growth curve of human KL mutant PDX sample implanted into nude mice treated with vehicle (n=5) or AHA-1031 (n=, right). ^∗^: p < 0.05, ^∗∗^: p < 0.01, ^∗∗∗^: p < 0.001, ^∗∗∗∗^: p < 0.0001.

We then performed additional in vivo studies with our lead candidate AHA-1031 because of the superior efficacy to other candidates. In the A549 xenograft model we also noted significantly decreased tumor growth in AHA-1031 treated animals compared to vehicle control. We however did not see any complete responders in this experiment (Figure 4C). This is likely because H2030 expresses higher levels of cell surface MICA/B resulting in increased anti-tumor activity of NK cells. We importantly did not see any observable toxicity in this model through measurement of bodyweight. To determine the immune-modulatory effects of our therapeutic we removed tumors from our A549 control and AHA-1031 treated animals and performed flow cytometry analysis. Compared to control treated animals, we found significantly increased lymphocyte infiltration (CD45/Live) and NK cell infiltration (NK/CD45). We also observed significantly increased activated NK cell populations through expression of key cytotoxic markers such as CD107a, Granzyme B, and Ki67 (Figure 4D). We also performed additional toxicity studies and incubated human PBMCs with increasing doses of AHA-1031. We found that AHA-1031 did not result in a significant increase in levels of pro-inflammatory cytokines IFNγ, IL-2, IL-4, and IL-17A (Supplementary Figure 4D)

To determine next whether our therapeutic is effective in a clinically relevant patient derived xenograft (PDX) model, we used a KRAS G12A LKB1 mutant NSCLC PDX in nude mice. We first removed untreated tumors and profiled MICA/B expression through flow cytometry and found cell surface expression of MICA/B (Figure 5E). Surface MICA/B was further stabilized, and levels increased with increasing doses of the AHA-1031. After confirmation of functionality, we implanted mice again with tumors and treated with either vehicle or AHA-1031. We observed a significant reduction in tumor growth with our therapeutic antibody compared to vehicle control (Figure 5F).

Importantly, our PDX results along with our A549 and H2030 models, demonstrate that AHA-1031 is effective across a range of MICA/B surface expression with H2030 demonstrating the highest expression and our KL PDX the lowest. Importantly similar stabilization of MICA in the KL PDX model and H2030 tumors were observed. This specific model we used is representative of KL mutant NSCLC harboring loss of LKB1 and a mutation in KRAS G12A. KRAS mutations are one of the most common oncogenic driver mutations of all cancers and specifically NSCLC, with G12C and G12D alternations as the most common. However, G12A mutations are also prevalent and found in around 7% of cases (28). Our therapeutic strategy therefore provides an alternative treatment to KRAS targeting through the cytotoxic effects of NK cells.

## Discussion/Conclusion

Here we show that our novel NK cell activating antibody, AHA-1031, shows potent anti-tumor activity and prevents MICA/B ligand shedding in preclinical models of KL mutant NSCLC. We first demonstrate that KL mutant NSCLC cell lines have high cell surface expression of MICA/B, and these ligands are secreted and shed in vitro. NSCLC patient blood sample analysis additionally shows that soluble MICA/B are found at very low levels in healthy people, however in NSCLC patients there is a significant increase in soluble MICA/B. Our antibody shows strong binding properties and specificity preventing MICA/B shedding across several solid tumor lines including lung, prostate, and pancreas. In vivo, we demonstrate in two KL mutant NSCLC xenograft models as well as a KL mutant PDX model that treatment with AHA-1031 monotherapy significantly inhibits tumor growth compared to vehicle treated animals. In the tumor tissues of treated mice, significantly increased immune cell infiltrates and activated NK cell populations are observed. Importantly we did not observe any measurable toxicity with our novel antibody.

KL mutant tumors have limited response to current immune activating therapies (29). Multiple studies have shown that compared to K driven tumors, KL mutant tumors have far worse response to ICB therapy, which is currently standard of care (18, 21). Interestingly, a recent study showed that LKB1 was one of the most acquired mutations after resistance to ICB therapy, noting its potential role as a resistance mechanism (30). While we do not propose that this treatment is uniquely specific to KL tumors, since KL tumors do not benefit from other therapies, NK cell modulating agents could be of significant benefit in this subtype.

To date, NK cell mediated therapies have lagged T-cell therapies across cancer types. Currently, the main therapeutics activating NK cells in clinical trials include cytokine stimulation such as IL-2 to increase NK cell activation and monoclonal antibodies such as Monalizumab which binds to NK cell inhibitory receptor NKG2A. Chimeric antigen receptor (CAR) NK cells have also been tested but have faced similar issues in solid tumors with immune cells not able to infiltrate the tumor (31). NK cells are particularly important in the lung and comprise 10-20% of all lymphocytes (32). In preclinical models of NSCLC, it was demonstrated that with progression of KRAS driven NSCLC there was a significant decrease in NK cell numbers over time. However, our treatment can overcome this by recruiting and activating NK cells. Currently, the only other anti-MICA antibody in clinical trials is CLN-619 (NCT05117476), which utilizes a WT human Ig1. However, our study demonstrates that substituting the WT Fc with an engineered IgG1-Fc significantly enhances NK cell cytotoxicity.

In conclusion, we show in our study that preventing MICA/B shedding is an effective therapeutic strategy in preclinical mouse models of KL mutant NSCLC. Importantly we show that our MICA antibody monotherapy is effective even at physiological levels of MICA expression. MICA/B are widely expressed across NSCLC and other cancer types making our therapeutic strategy widely applicable.

## Acknowledgements

EAA is a Cancer Prevention and Research Institute of Texas (CPRIT) Scholar in Cancer Research. EAA was supported by CPRIT Scholar Award RR160080, NIH (National Institutes of Health)-R01CA276058, 1R01CA289500-01, Department of Defense (DOD) W81XWH-21-1-0856, American Cancer Society RSG-22-051-01-IBCD and William Guy Forbeck Foundation. EAA was also supported by the NCCN Foundation®. Any opinions, findings, and conclusions expressed in this material are those of the author(s) and do not necessarily reflect those of National Comprehensive Cancer Network® (NCCN®) or the NCCN Foundation. JDM, DG, and EAA were supported by NIH 5P50CA070907. RRK was supported by 5T32CA124334. We thank Simmons Cancer Center for Cancer Center resources (P30CA142543).

## Author Contributions

Conceptualization: EAA, HB. Methodology and analysis: RRK, MS, LTG, QD, LG, YN, KM, SK. Writing and editing: RRK, MS, HB, EAA. Funding acquisition: EAA, HB. Resources: EAA, HB, ZLL, JDM, DEG.

## Conflict of Interest Disclosure

H.B, M.S are employees of Aakha Biologics and have shares in the company. L.T.G was an employee of Aakha Biologics and has shares in the company. Z.L.L is an employee of Alloy Therapeutics. H.B., M.S., Z.L.L., and L.T.G have shares in Alloy. H.B, M.S, Z.L.L and L.T.G are co-inventors of antibody-related patent. D.E.G has research funding from Astra-Zeneca, BerGenBio, Karyopharm, and Novocure; stock ownership from Gilead, Medtronic, and Walgreens; consulting/advisory Boards of Astra-Zeneca, Catalyst Pharmaceuticals, Daiichi-Sankyo, Elevation Oncology, Janssen Scientific Affairs, LLC, Jazz Pharmaceuticals, Regeneron Pharmaceuticals, and Sanofi; intellectual property (U.S. patent 11,747,345; U.S. patent applications 17/045,482, 63/386,387, 63/382,972, 63/382,257; co-founder and Chief Medical Officer, OncoSeer Diagnostics, Inc. JDM receives licensing fees for cell lines. Other authors declare no relevant conflicts of interest.

---

**Supplemental Figure 1:**
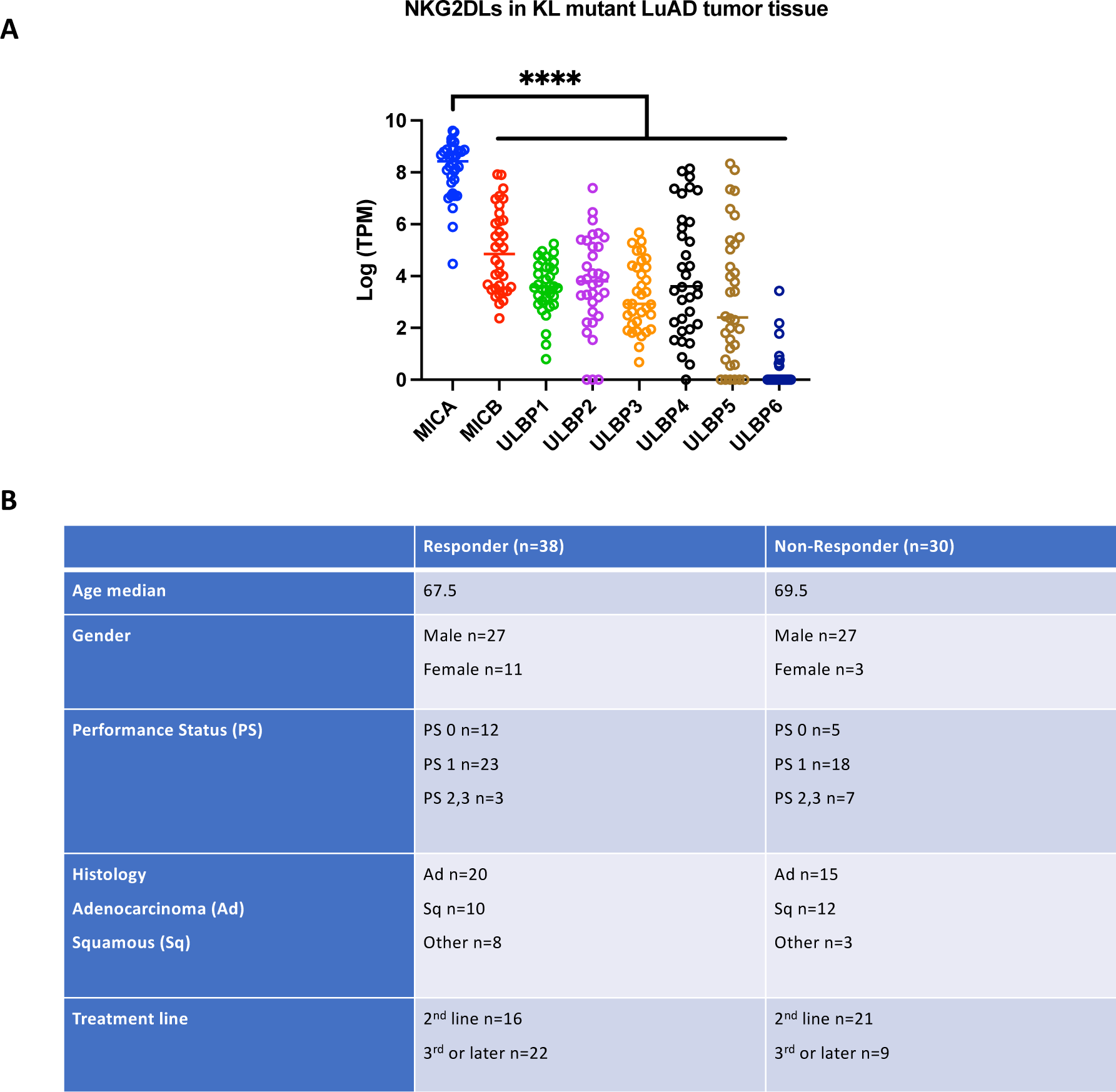
Patient information. (A) Expression of NKG2DLs specifically in KL mutant lung adenocarcinomas (TCGA). (B) Clinical and demographic details of patients whose blood samples were analyzed in Figure

**Supplemental figure 2:**
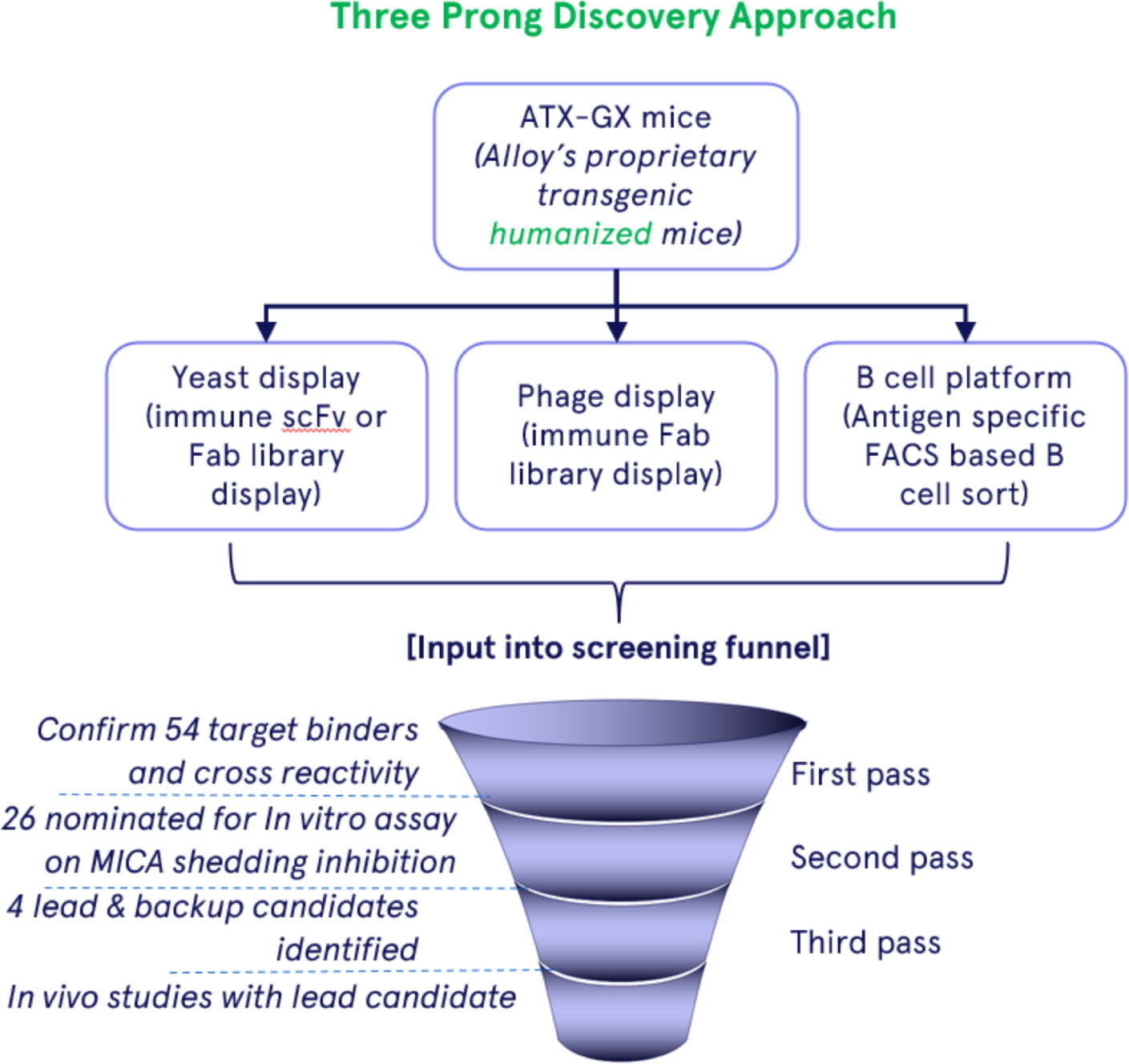
Identification of novel anti-MICA antibodies. Diagram illustrating Alloy Therapeutics’ (ATX-Gx™) proprietary transgenic mouse platform used to generate fully human anti-MICA/B antibodies with reduced immunogenicity risk. A summary of the discovery process shows numerous hits were identified and 54 antibodies were selected based on sequence diversity and initial affinity measurements. These antibodies were then produced as full-length IgG1 antibodies. Subsequently, a series of biophysical, developability (drug-like properties), and in vitro assays were conducted to identify the first-generation lead candidate, AHA-1032

**Supplemental Figure 3:**
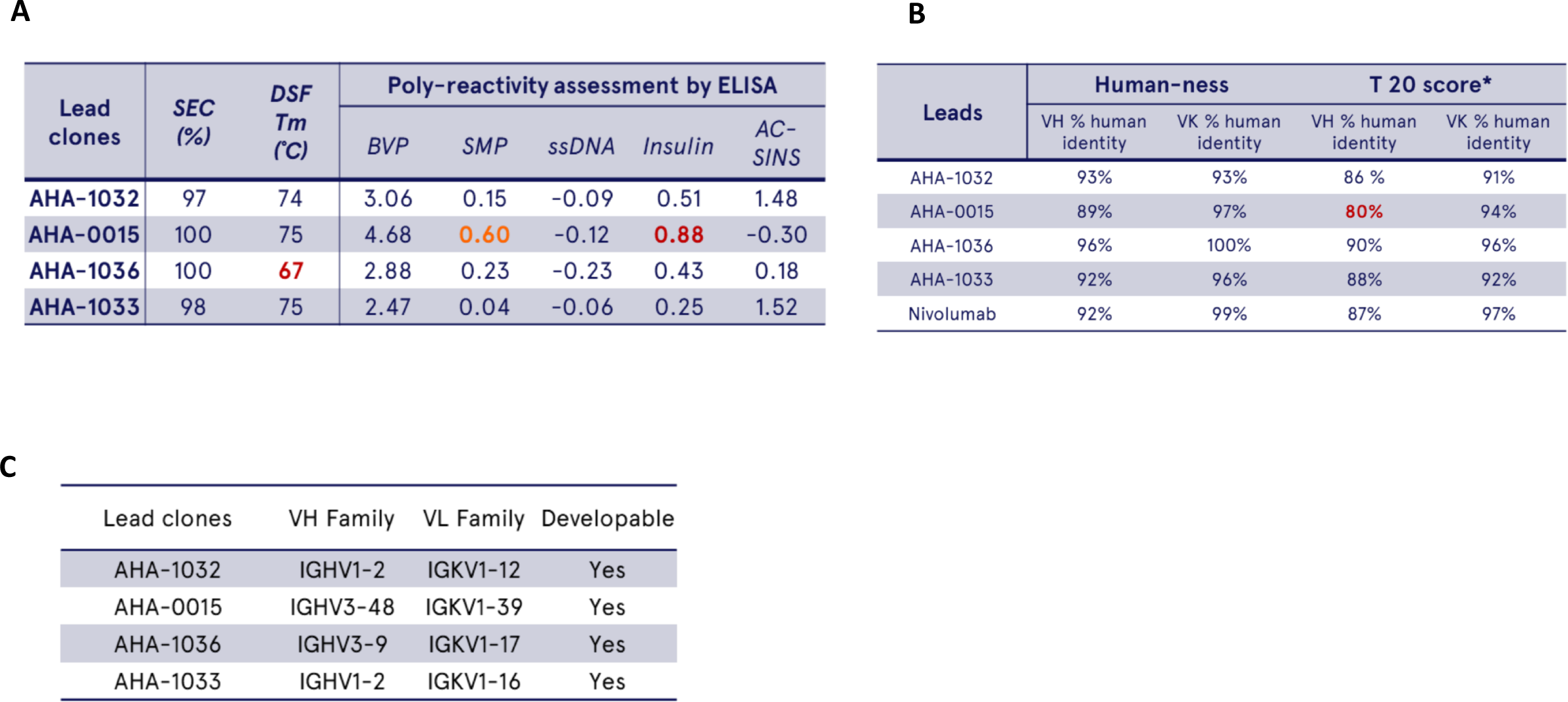
Specifications of anti-MICA antibodies. (A) Physicochemical Properties of Alpha3-Specific Lead Antibodies (Table A) Size Exclusion Chromatography (SEC) and nano-Differential Scanning Fluorimetry (nanoDSF) Analysis. The percentage of the main peak by SEC following Protein A purification was measured. The melting temperature (Tm) of the lead candidates was determined using nanoDSF. All lead candidates were assessed for non-specific binding to polyreactive reagents using ELISA. (B) Development and Humanization of Anti-MICA Antibodies. Anti-MICA antibodies were generated from human transgenic mice and the degree of humanness for all lead antibodies was evaluated using IMGT database. The humanness score for each lead candidate was calculated, and the T20 score was reported. The T20 score ranges from 0 to 100, with higher scores indicating a more human-like antibody. Full-length sequences scoring above 80 are generally considered human-like. (C) Germline Analysis and Developability Assessment of Lead Candidates. The variable regions of all lead candidates were analyzed using the IMGT database to determine germline origins. The corresponding degree of developability for each candidate was evaluated.

**Supplemental Figure 4:**
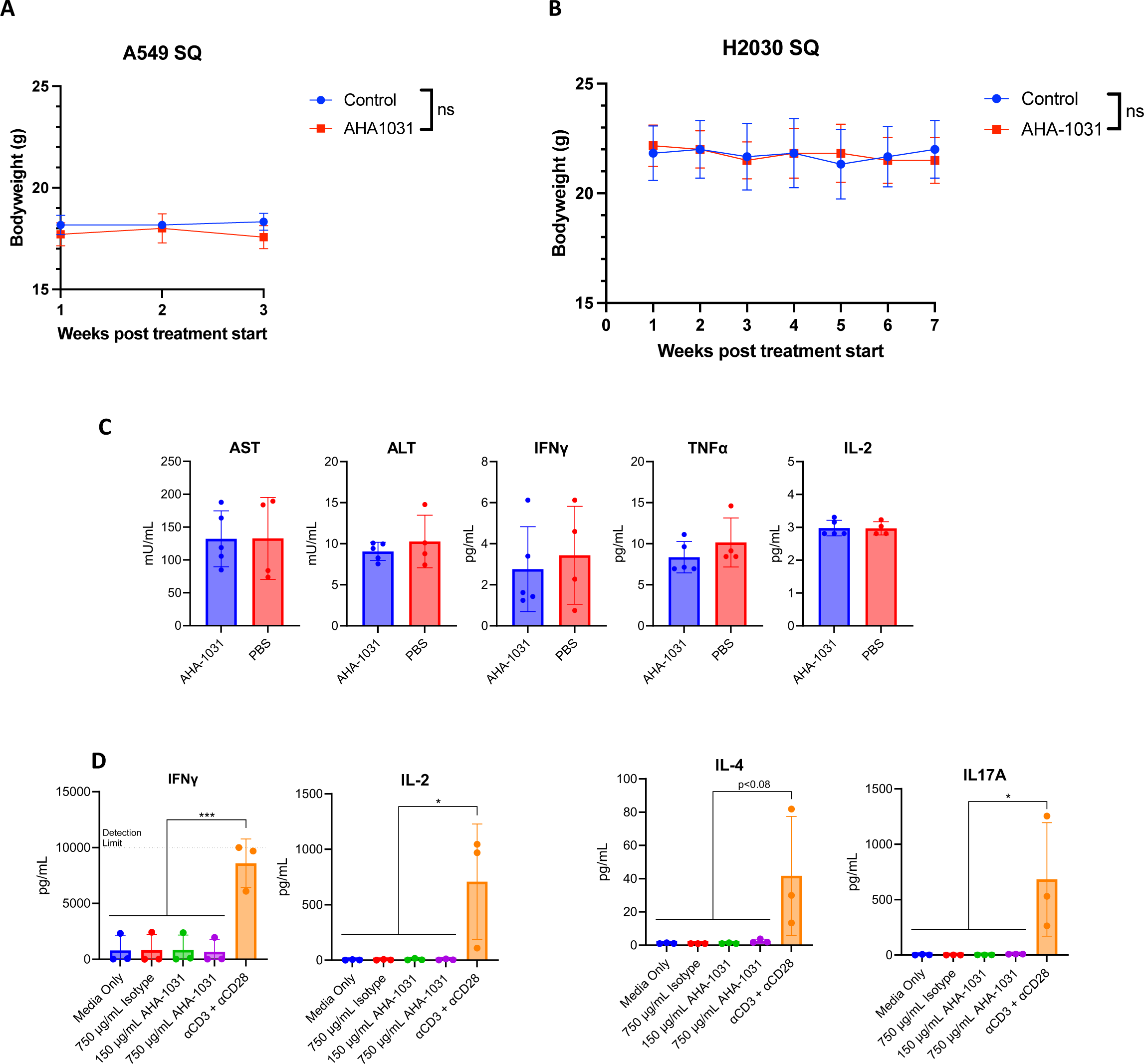
Toxicity assessment of MICA antibody. (A) Bodyweight measurements of A549 subcutaneously implanted mice measured at 1-, 2-, and 3-weeks post treatment start in control (n=4) or AHA-1031 (n=4) treated mice (B) Bodyweight measurements of H2030 subcutaneously implanted mice measured at 1-7 weeks post treatment start in control (n=6) or AHA-1031 (n=6) treated mice (C) Liver toxicity analysis of serum of B16F10-MICA bearing mice treated with repeated doses of AHA-1031 (n=5) or vehicle (n=4) (D) Cytokine release assay of donor human PBMC’s treated with vehicle, increasing doses of AHA-1031, or anti-CD3-CD28 positive control as detected through ELISA

## REFERENCES

1. Cong J, Wang X, Zheng X, Wang D, Fu B, Sun R, et al. Dysfunction of Natural Killer Cells by FBP1-Induced Inhibition of Glycolysis during Lung Cancer Progression. Cell Metab. 2018;28(2):243–55 e5.

2. Myers JA, Schirm D, Bendzick L, Hopps R, Selleck C, Hinderlie P, et al. Balanced engagement of activating and inhibitory receptors mitigates human NK cell exhaustion. JCI Insight. 2022;7(15).

3. Eleme K, Taner SB, Onfelt B, Collinson LM, McCann FE, Chalupny NJ, et al. Cell surface organization of stress-inducible proteins ULBP and MICA that stimulate human NK cells and T cells via NKG2D. J Exp Med. 2004;199(7):1005–10.

4. Xing S, Ferrari de Andrade L. NKG2D and MICA/B shedding: a ‘tag game’ between NK cells and malignant cells. Clin Transl Immunology. 2020;9(12):e1230.

5. Ferrari de Andrade L, Tay RE, Pan D, Luoma AM, Ito Y, Badrinath S, et al. Antibody-mediated inhibition of MICA and MICB shedding promotes NK cell-driven tumor immunity. Science. 2018;359(6383):1537-42.

6. Chen Y, Lin G, Guo ZQ, Zhou ZF, He ZY, Ye YB. Effects of MICA expression on the prognosis of advanced non-small cell lung cancer and the efficacy of CIK therapy. PLoS One. 2013;8(7):e69044.

7. Zhu M, Huang Y, Bender ME, Girard L, Kollipara R, Eglenen-Polat B, et al. Evasion of Innate Immunity Contributes to Small Cell Lung Cancer Progression and Metastasis. Cancer Res. 2021;81(7):1813–26.

8. Zhao Y, Chen N, Yu Y, Zhou L, Niu C, Liu Y, et al. Prognostic value of MICA/B in cancers: a systematic review and meta-analysis. Oncotarget. 2017;8(56):96384–95.

9. Waldhauer I, Goehlsdorf D, Gieseke F, Weinschenk T, Wittenbrink M, Ludwig A, et al. Tumor-associated MICA is shed by ADAM proteases. Cancer Res. 2008;68(15):6368–76.

10. Zingoni A, Vulpis E, Loconte L, Santoni A. NKG2D Ligand Shedding in Response to Stress: Role of ADAM10. Front Immunol. 2020;11:447.

11. Rebmann V, Schutt P, Brandhorst D, Opalka B, Moritz T, Nowrousian MR, et al. Soluble MICA as an independent prognostic factor for the overall survival and progression-free survival of multiple myeloma patients. Clin Immunol. 2007;123(1):114–20.

12. Cascone R, Carlucci A, Pierdiluca M, Santini M, Fiorelli A. Prognostic value of soluble major histocompatibility complex class I polypeptide-related sequence A in non-small-cell lung cancer - significance and development. Lung Cancer (Auckl). 2017;8:161–7.

13. de Castro G, Jr., Kudaba I, Wu YL, Lopes G, Kowalski DM, Turna HZ, et al. Five-Year Outcomes With Pembrolizumab Versus Chemotherapy as First-Line Therapy in Patients With Non-Small-Cell Lung Cancer and Programmed Death Ligand-1 Tumor Proportion Score >/= 1% in the KEYNOTE-042 Study. J Clin Oncol. 2023;41(11):1986–91.

14. Chevallier M, Borgeaud M, Addeo A, Friedlaender A. Oncogenic driver mutations in non-small cell lung cancer: Past, present and future. World J Clin Oncol. 2021;12(4):217–37.

15. Govindan R, Ding L, Griffith M, Subramanian J, Dees ND, Kanchi KL, et al. Genomic landscape of non-small cell lung cancer in smokers and never-smokers. Cell. 2012;150(6):1121–34.

16. Krishnamurthy N, Goodman AM, Barkauskas DA, Kurzrock R. STK11 alterations in the pan-cancer setting: prognostic and therapeutic implications. Eur J Cancer. 2021;148:215–29.

17. Rosellini P, Amintas S, Caumont C, Veillon R, Galland-Girodet S, Cuguilliere A, et al. Clinical impact of STK11 mutation in advanced-stage non-small cell lung cancer. Eur J Cancer. 2022;172:85–95.

18. Skoulidis F, Goldberg ME, Greenawalt DM, Hellmann MD, Awad MM, Gainor JF, et al. STK11/LKB1 Mutations and PD-1 Inhibitor Resistance in KRAS-Mutant Lung Adenocarcinoma. Cancer Discov. 2018;8(7):822–35.

19. McMillan EA, Ryu MJ, Diep CH, Mendiratta S, Clemenceau JR, Vaden RM, et al. Chemistry-First Approach for Nomination of Personalized Treatment in Lung Cancer. Cell. 2018;173(4):864–78 e29.

20. Cancer Genome Atlas Research N. Comprehensive molecular profiling of lung adenocarcinoma. Nature. 2014;511(7511):543-50.

21. Alvez MB, Edfors F, von Feilitzen K, Zwahlen M, Mardinoglu A, Edqvist PH, et al. Next generation pan-cancer blood proteome profiling using proximity extension assay. Nat Commun. 2023;14(1):4308.

22. Pan M, Wang F, Nan L, Yang S, Qi J, Xie J, et al. alphaVEGFR2-MICA fusion antibodies enhance immunotherapy effect and synergize with PD-1 blockade. Cancer Immunol Immunother. 2023;72(4):969–84.

23. Deng J, Thennavan A, Dolgalev I, Chen T, Li J, Marzio A, et al. ULK1 inhibition overcomes compromised antigen presentation and restores antitumor immunity in LKB1 mutant lung cancer. Nat Cancer. 2021;2(5):503–14.

24. Best SA, Hess JB, Souza-Fonseca-Guimaraes F, Cursons J, Kersbergen A, Dong X, et al. Harnessing Natural Killer Immunity in Metastatic SCLC. J Thorac Oncol. 2020;15(9):1507–21.

25. Toledo-Stuardo K, Ribeiro CH, Canals A, Morales M, Garate V, Rodriguez-Siza J, et al. Major Histocompatibility Complex Class I-Related Chain A (MICA) Allelic Variants Associate With Susceptibility and Prognosis of Gastric Cancer. Front Immunol. 2021;12:645528.

26. Wu JD, Higgins LM, Steinle A, Cosman D, Haugk K, Plymate SR. Prevalent expression of the immunostimulatory MHC class I chain-related molecule is counteracted by shedding in prostate cancer. J Clin Invest. 2004;114(4):560–8.

27. Park JE, Kim SE, Keam B, Park HR, Kim S, Kim M, et al. Anti-tumor effects of NK cells and anti-PD-L1 antibody with antibody-dependent cellular cytotoxicity in PD-L1-positive cancer cell lines. J Immunother Cancer. 2020;8(2).

28. Zhao D, Li H, Mambetsariev I, Mirzapoiazova T, Chen C, Fricke J, et al. Clinical and Molecular Features of KRAS-Mutated Lung Cancer Patients Treated with Immune Checkpoint Inhibitors. Cancers (Basel). 2022;14(19).

29. Jeanson A, Tomasini P, Souquet-Bressand M, Brandone N, Boucekine M, Grangeon M, et al. Efficacy of Immune Checkpoint Inhibitors in KRAS-Mutant Non-Small Cell Lung Cancer (NSCLC). J Thorac Oncol. 2019;14(6):1095–101.

30. Ricciuti B, Arbour KC, Lin JJ, Vajdi A, Vokes N, Hong L, et al. Diminished Efficacy of Programmed Death-(Ligand)1 Inhibition in STK11- and KEAP1-Mutant Lung Adenocarcinoma Is Affected by KRAS Mutation Status. J Thorac Oncol. 2022;17(3):399–410.

31. Liu S, Galat V, Galat Y, Lee YKA, Wainwright D, Wu J. NK cell-based cancer immunotherapy: from basic biology to clinical development. J Hematol Oncol. 2021;14(1):7.

32. Cong J, Wei H. Natural Killer Cells in the Lungs. Front Immunol. 2019;10:1416.

